# Genetic variance and indirect genetic effects for affiliative social behavior in a wild primate

**DOI:** 10.1101/2022.08.29.505695

**Authors:** Emily M. McLean, Jacob A. Moorad, Jenny Tung, Elizabeth A. Archie, Susan C. Alberts

## Abstract

Affiliative social behaviors are linked to fitness components in multiple species. However, the role of genetic variance in shaping affiliative social behaviors remains largely unknown, limiting our understanding of how these behaviors can respond to natural selection. Here, we employed the ‘animal model’ to estimate both environmental and genetic sources of variance and covariance in grooming behavior in the well-studied Amboseli baboon population in Kenya. We found that grooming given, grooming received, and total grooming all are similarly heritable (h^2^=0.22, h^2^=0.16, and h^2^=0.26 respectively), and that rank and the presence of kin contribute to environmental variance. We detected small but measurable indirect genetic effects of partner identity on the amount of grooming given within dyadic grooming partnerships. The genetic correlation between grooming given and grooming received was exceptionally strong and positive (R=0.94 ± 0.12), and the indirect and direct genetic effects for grooming given were also strongly positively correlated (R=0.86 ± 0.06). Our results provide insight into the evolvability of affiliative behavior in wild animals, including the possibility for correlations between direct and indirect genetic effects to accelerate the response to selection. As such they provide novel information about the genetic architecture of social behavior in nature, with important implications for the evolution of cooperation and reciprocity.

## INTRODUCTION

In humans and many other social mammals, the frequency and intensity of affiliative interactions with others (‘social integration’) predict individual survival and reproductive success (e.g., Holt-Lunstad et al. 2010; Stanton and Mann 2012; McFarland and Majolo 2013; Vander Wal et al. 2015; Ellis et al. 2017; Thompson and Cords 2018, Cameron et al. 2009; Schülke et al. 2010, Feldblum et al. 2021). Strong, differentiated affiliative relationships confer several potential fitness benefits to group living animals, including parasite removal (Ezenwa et al. 2016), access to mating opportunities (Diaz-Muñoz et al. 2014), decreased intra-group conflict (Silk 2002) and enhanced success in within and between group competitive encounters (Wrangham 1980).

Given the links between social relationships and fitness in highly social species, natural selection likely favors more socially integrated individuals. However, despite the clear and compelling links between social relationships and fitness, we have a limited understanding of how differentiated social relationships evolve. Specifically, for an evolutionary response to selection to occur, phenotypic variation in social integration must have an underlying heritable component. Evidence from human populations suggests that loneliness and social integration are weakly to modestly heritable and subject to strong environmental effects (Day et al. 2018, Abdellaoui et al. 2019), including indirect genetic effects (IGEs: i.e., the effects of other genotypes present in the environment; sometimes termed “social genetic effects”). However, in wild, non-human vertebrate populations, few studies have directly investigated the heritability of affiliative social behaviors (but see Blomquist and Brent, 2014). Instead, studies to date have focused on social network metrics (e.g., Fowler et al. 2009; Lea et al. 2010; Brent et al. 2012) or cooperative behaviors (e.g., Bleakley and Brodie 2009, Kasper et al. 2017, Houslay et al. 2021). Measurable heritability values for both types of traits suggest possible pathways through which social integration could affect fitness (e.g., cooperation can emerge from social bonds: Berghänel et al. 2011). However, because social network metrics are not traits that can be measured for single individuals, and because cooperative behavior differs from social affiliation *per se*, direct estimates of the heritability of social affiliation remain lacking.

Taken together, these studies highlight the need for further investigation into the genetic architecture of social affiliation, including the relative contributions of genetic and environmental variation to phenotypic variation in wild populations. Here, we contribute to filling this gap by combining detailed, long-term data on individual social relationships with the extensive pedigree available for the well-studied Amboseli baboon population. Importantly, these datasets allow us to investigate genetic variance and covariance in affiliative behaviors at the level of the individual, as well as indirect genetic effects for social affiliation at the level of the dyadic social relationship.

### Genetic covariances, indirect genetic effects, and the evolution of social behavior

When two traits are genetically correlated (i.e., exhibit genetic covariance caused by linkage or pleiotropy), the evolutionary response to selection may be constrained or accelerated, depending on the direction of the covariance and the direction of selection (Lande and Arnold 1983). This occurs because genetic correlations can prevent the independent evolution of a single trait. For instance, sexual attractiveness in male guppies (*Poecilia reticulata*) is heritable but negatively genetically correlated with offspring survival (Brooks 2000). Consequently, positive selection to increase male attractiveness is constrained, because the genetic change required to respond to selection also tends to reduce offspring survival, creating a trade-off that arises at the genetic level.

Furthermore, in a social environment, genetic effects on phenotype are not limited to an individual’s own genes (direct genetic effects), but potentially include the genes of its social partners (indirect genetic effects). Selection can act on indirect genetic effects whether social partners are related (Wolf et al. 1998) or unrelated, particularly when IGEs are correlated with the direct genetic effects (DGEs) of an individual’s genotype on its own phenotype. For instance, Wilson et al. (2009) found a positive genetic correlation between DGEs and IGEs for some aggressive phenotypes in a lab population of deer mice (*Peromyscus maniculatus*), implying that the same genotypes that promote aggression in the bearer also promote aggression in those with whom it interacts. Selection for increased aggression, then, would result in evolution of the social environment as well a change in frequency of ‘aggressive alleles’: each successive generation would experience a more aggressive social environment than that of their parents (even the individuals who themselves did not carry ‘aggressive alleles’) and hence would themselves be more aggressive. That is, phenotypic evolution would be greater than expected if DGEs and IGEs were independent (for a fuller treatment of the quantitative genetic approach to understanding indirect genetic effects, see Moore et al. 1997; Wolf et al. 1998; Hunt and Simmons 2002; Bijma and Wade 2008 and references therein).

Evidence for the importance of indirect genetic effects in explaining variation in social environment-associated traits is accumulating (Baud et al. 2017, Kong et al. 2018, Mostafavi et al. 2020). However, little work has focused on IGEs for behaviors that contribute to differentiated, dyadic affiliative relationships, especially in wild animal populations. Here we contribute to the limited knowledge in this area by investigating the heritability and genetic architecture (including IGEs) of social grooming, a common affiliative behavior in primates with known links to the survival components of fitness (Silk et al. 2003, Silk et al. 2010, Archie et al. 2014, Campos et al. 2020).

### Grooming behavior in non-human primates

Grooming is a primary means by which many non-human primates establish and maintain differentiated, affiliative social bonds (Silk 1987). While grooming relationships are common among kin and mating pairs, strong and enduring bonds also occur between unrelated pairs (Silk 1987). Grooming involves manually picking through and cleaning the fur of debris and ectoparasites and is known to reduce disease risk (Tanaka and Takefushi 1993; Sánchez-Villagra et al. 1998; Akinyi et al. 2013). However, grooming is common even when ectoparasites are eliminated (e.g., in captive primates), and the importance of grooming for social bonding in primates is widely recognized (Dunbar 1991; Silk 2007; Cords 2012). Social grooming can reduce tension and aggression between individuals (e.g., Saunders and Hausfater 1988), and in some wild populations, social grooming can occupy as much as 20% of an animal’s time budget (Dunbar 1991).

In a number of primate species, grooming relationships are generally reciprocal: within dyads, individuals who give more grooming also receive more grooming (e.g., see meta-analysis in Schino and Aureli 2008; also chimpanzees: Gomes et al. 2009; capuchins: Schino et al. 2009; baboons: Silk and Frank 2009; Silk et al. 2010). In baboons, the most enduring social relationships (those that last years rather than months), tend to be highly reciprocal or ‘equitable’ (Silk et al. 2006a, 2010). Thus, the grooming an individual receives, and the grooming they give to others are strongly phenotypically correlated, even though these phenotypes may have opposing fitness consequences for an individual animal (see Keverne et al. 1989, Wittig et al. 2008, Akinyi et al. 2013, Young et al. 2014 for benefits of receiving grooming and Dunbar and Sharman 1988, Schino 2007 for the small cost of giving grooming). Importantly, females appear to make decisions about who to groom based partly on the grooming behavior of their social partners (Schino 2007; Schino and Aureli 2008). Thus, if grooming behavior is shaped by an individual’s genotype, it follows that giving and receiving grooming are strong candidates for behaviors that are influenced by indirect genetic effects.

### Goals of this analysis

Here, we use data on >100,000 grooming interactions (n=224 baboons) collected from the well-studied baboons of the Amboseli region of Kenya to pursue three goals (Alberts and Altmann 2012, Alberts 2019). First, we describe how grooming behavior responds to social and non-social environmental effects. Second, we estimate variance explained by genetic effects (both direct and indirect) on grooming. Third, we measure the genetic relationship between grooming given and grooming received. We explicitly differentiate between direct and indirect genetic effects on grooming behavior, to better understand how these affiliative behaviors might respond to selection.

We address all three goals by employing the ‘animal model’, a mixed effects linear model that estimates both environmental and genetic sources of variance and covariance in phenotypes (see Methods, also Lynch and Walsh 1998; Kruuk 2004). We consider two suites of grooming phenotypes: (i) *aggregate* measures of grooming for each adult female (i.e., all of the grooming that an adult female gives to and/or receives from all other adult females, regardless of partner identity), and (ii) a *dyadic* measure of grooming (i.e., grooming given by an adult female to a specific adult female grooming partner, summarized in a yearly index). With our aggregate measures (Figure 1A), we investigated environmental and direct genetic sources of variance and the genetic covariance between grooming given and grooming received. With our dyadic measure (Figure 1B), we investigated environmental, direct, and indirect genetic sources of variance, as well as the genetic covariance between direct and indirect genetic variance.

**Figure 1.**
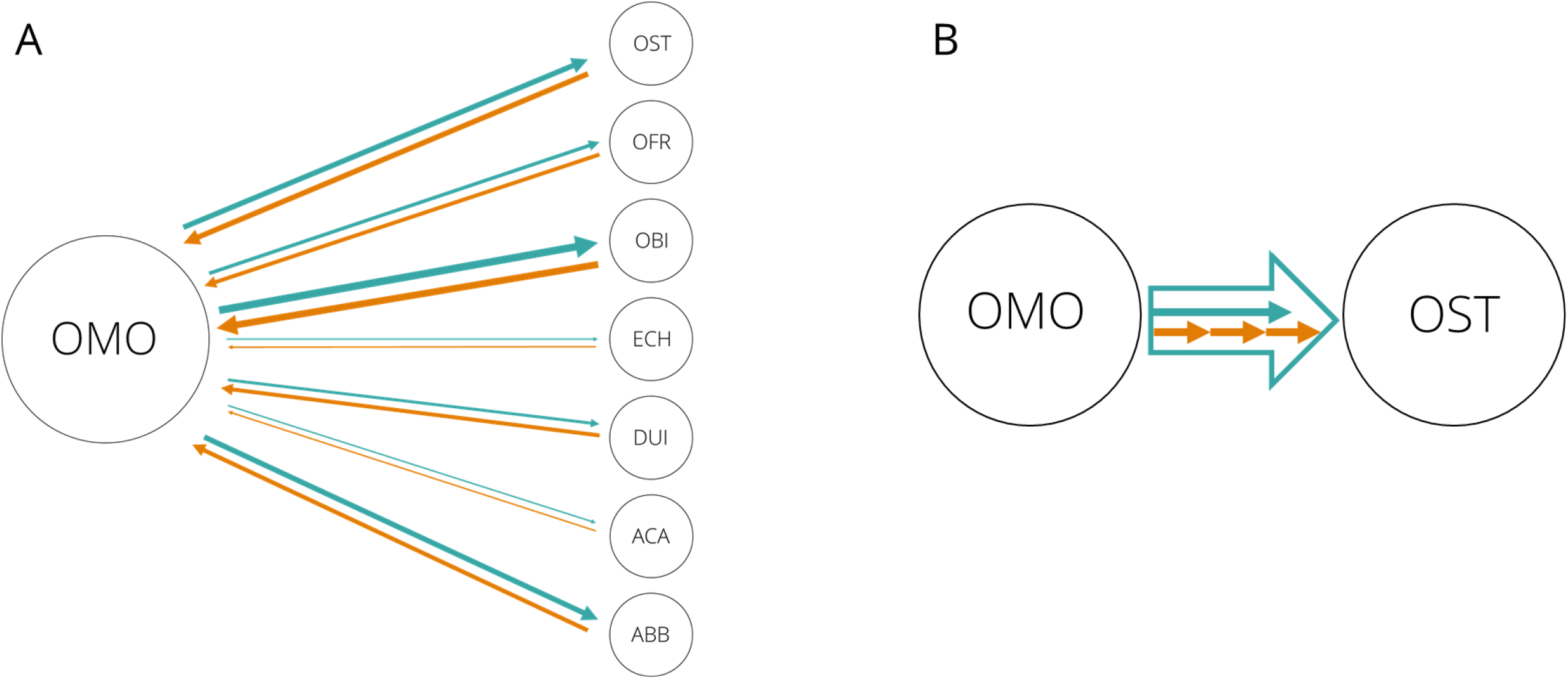
Visualization of grooming metrics. (A) An example of our aggregate grooming indices, representing grooming given to and received by adult female baboon OMO in 2003, considering all other adult females in her social group that year (individual females are identified by 3-letter codes). Green arrows represent grooming given by OMO; orange arrows represent grooming received by OMO. The width of the arrow indicates the relative frequency of grooming observed; wider arrows indicate higher relative frequencies. OMO’s aggregate grooming given for 2003 is the sum of the green arrows, her aggregate grooming received is the sum of the orange arrows, and her total grooming is the sum of those two metrics. (B) An example of our dyadic index. Here, we observed the grooming given by OMO to adult female OST in a single year, represented by the large, green block arrow. This phenotype is shaped by two components: the tendency of OMO to give grooming (direct genetic effects), represented by the small green arrow, and the tendency of OST to elicit grooming from her partners (indirect genetic effects) represented by the dashed orange arrow.

## METHODS

### Study population and grooming data collection

The Amboseli baboon population of southern Kenya has been the subject of ongoing research for five decades (Alberts and Altmann 2012; Alberts 2019). The ancestry of baboons in this population is primarily yellow baboon, but all individuals contain low to moderate levels of anubis baboon ancestry due to naturally occurring admixture with neighboring anubis populations (Alberts and Altmann 2001; Vilgalys et al. 2022). All animals in the social groups under study (the ‘study groups’) are individually recognized on sight based on unique morphological and facial features. All demographic and life-history events (births, maturation events, immigrations, deaths, and emigrations) are recorded as part of the near-daily monitoring of the study groups.

Our grooming data consisted of counts of grooming events between adult females, with both the giver and receiver of grooming recorded. Grooming was recorded whenever one animal used both hands to pick through the fur of a second animal. Only grooming events between adult females were considered for this analysis. We collected grooming counts during systematic monitoring of the population, following a sampling protocol that is designed to avoid potential biases that could result from uneven sampling of study subjects (see Supplementary Methods).

Our study subjects were all adult female baboons (N=224) present in the study groups between January 1983 and June 2017 for whom we have known pedigree links and enough genetic material to calculate their anubis-yellow ‘admixture score’ (see Tung et al 2008). Females were considered adults if they had attained menarche. The resulting dataset represented 115,149 grooming interactions collected during 1,868 female-years of life, with a median of 400.5 interactions per individual.

The research in this study was approved by the Institutional Animal Care and Use Committee (IACUC) at Duke University (no. A273-17-12) and adhered to the laws and guidelines of the Kenyan government.

### Grooming indices

#### Aggregate indices of grooming

To determine the direct genetic effect variances and covariance of grooming given and grooming received, we used the counts of grooming bouts between adult females to calculate a set of aggregate grooming indices, adjusted for observer effort (see Supplementary Methods). For each adult female in each year of her adult life we calculated an index of *aggregate grooming given* that reflects the frequency with which she groomed other adult females, relative to the grooming given by all other adult females alive in the same year. Similarly, we calculated an index of *aggregate grooming received* that reflects the relative frequency with which she received grooming from other adult females (Figure 1A; see also Supplementary Methods, Figure S1A,B and Archie et al. 2014). Positive values for either index indicate females with relatively high frequencies of grooming (given or received, respectively) for females in the population in that year; negative values represent females with relatively low frequencies of grooming for that year. Our *aggregate index of total grooming* is the average of the *aggregate grooming given* and *aggregate grooming received* indices, calculated for each adult female for each year of her adult life (see Supplementary Methods and Figure S1C).

To determine the phenotypic correlation between our repeated measures of aggregate grooming given and aggregate grooming received we used the ‘rmcorr’ package in R v3.4.1 (Bakdash and Marusich 2017). We calculated this correlation using all of our data points (n=224 females, n=1,868 female years).

#### Dyadic index of grooming

In order to measure indirect genetic effects on grooming we calculated a yearly *dyadic grooming index* for each pair of adult females that were co-resident in a social group for at least 60 days during the calendar year and that had at least one grooming interaction (Figure 1B). The primary benefit of using the dyadic index was that it allowed us to investigate both direct genetic effects and the indirect genetic effects of social partners, as well as the correlation between thes effects. In contrast, the aggregate indices only allowed us to investigate direct genetic effects. We also used the dyadic grooming index to investigate environmental and direct genetic effects, which we expected to corroborate the results of our aggregate grooming measure.

Positive values of the dyadic index indicate cases in which an adult female gave high frequencies of grooming to a specific partner relative to all other partner pairs in the population for that year, while negative values indicate cases in which an adult female gave relatively low frequencies of grooming to a specific partner (see Supplementary Methods and Figure S1D for details).

#### The ‘animal model’ approach

To partition the phenotypic variance in these measures of social affiliation into additive genetic and other variance components we combined pedigree information and phenotypic values in a mixed effects model, the ‘animal model’ (see Lynch and Walsh 1998, Kruuk 2004). We constructed our pedigree based on long-term demographic records and on genetic parentage assignment carried out using 7-14 microsatellite genotypes. These procedures have become standard in the study population and have allowed us to produce a pedigree that includes more than 1,500 individuals (Galezo et al. 2022). The subset of this pedigree necessary to describe the relationships between all 224 of our study subjects consists of 308 individuals (see Supplementary Methods).

The animal model includes breeding value as a random effect. True breeding values are unknown, but they can be estimated based on the expected covariance in additive genetic effects between relatives (see Lynch and Walsh 1998, Kruuk 2004). The matrix form of the animal model can be represented by:

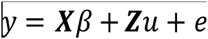

where *y* is the vector of phenotypic observations, *β* is the vector of fixed effects, *u* is the vector of random effects, **X** and **Z** are design matrices relating the fixed effects and random effects to each individual and *e* is the vector of residual errors. We discuss the robustness of this model to the pedigree structure of our population, grooming interactions between kin and admixture-related variation in genetic ancestry in the Supplementary Methods.

#### Goal 1: Fixed effects: Social and non-social influences on female grooming behavior

In our quantitative genetic animal models, we included fixed effects of other variables known or predicted to influence grooming behavior (see Table S1 for complete descriptions). These include (i) age, (ii) ordinal dominance rank, (iii) male and female group size, (iv) presence of mother, adult daughters, and adult maternal sisters, (v) measures of pedigree relatedness to other adult females in the group or to the dyadic partner, and (vi) individual admixture score (Table S1). The specific metrics we used to model these effects varied slightly according to whether we were analyzing aggregate measures or the dyadic index of grooming (Table S1, Table 1).

**Table 1.** Social and non-social environmental effects on aggregate and dyadic grooming indices.

|  | **Aggregate Grooming Given** | **Aggregate Grooming Received** | **Aggregate Total Grooming** | **Dyadic Grooming Index** | **Interpretation of effect size directions and estimates** |
| --- | --- | --- | --- | --- | --- |
| **Individual traits** |  |  |  |  |  |
| Focal dominance rank | **-0.01<sup>†</sup>**<br>**p=0.049** | -0.01<br>p=0.120 | **-0.02**<br>**p=0.031** | -0.035<br>p=0.20 | Aggregate indices: higher ranking females (those with lower numerical rank) gave more grooming, and engaged in more total grooming. The estimates represent the expected change in each grooming index in response to an increase of one numerical focal rank position.<br>Dyadic index: no effect. |
| Partner dominance rank | NA | NA | NA | -0.030<br>p=0.56 | Dyadic index: no effect. |
| Focal-partner rank interaction | NA | NA | NA | **0.002**<br>**p<0.0001** | Dyadic index: High ranking focals groomed high ranking partners more than low ranking partners. Low ranking focals groomed low ranking partners more than high ranking partners (Figure S3). |
| Focal age | **-0.09**<br>**p<0.0001** | **0.02**<br>**p<0.0001** | **-0.06**<br>**p<0.0001** | **-0.030**<br>**p<0.0001** | Aggregate indices: Older females received more grooming; younger females gave more grooming and engaged in more total grooming.<br>Dyadic index: Females gave less grooming to their partners when they (the focals) were older. The estimates represent the expected change in each grooming index in response to a one year change in focal age. |
| Partner age | NA | NA | NA | **0.006**<br>**p=0.023** | Dyadic index: Females gave more grooming to their partners when their partners were older. The estimate represents the expected change in the dyadic grooming index for a one year change in partner age. |
| Focal-partner age interaction | NA | NA | NA | -0.00003<br>p=0.953 | Dyadic index: no effect. |
| **Family effects** |  |  |  |  |  |
| Co-residency with mother | **0.14**<br>**p=0.017** | **0.22**<br>**p=0.0001** | **0.36**<br>**p=0.0004** | NA | Aggregate indices: Females who spent more of the time period co-residing with their mother gave more grooming, received more grooming, and groomed more in total. The estimate represents the expected difference in aggregate grooming indices between females that spent none of the time period with their mom and females that spent 100% of the time period with their mom. |
| Co-residency with adult daughters | **0.37**<br>**p<0.0001** | **0.24**<br>**p<0.0001** | **0.61**<br>**p<0.0001** | NA | Aggregate indices: Females who spent more of the time period co-residing with adult daughters gave more grooming, received more grooming, and groomed more in total. The estimate represents the expected difference in aggregate grooming indices between females that spent none of the time period with any adult daughters and females that spent 100% of the time period with an average of 1 adult daughter. |
| Co-residency with adult maternal sisters | -0.02<br>p=0.457 | **0.08**<br>**p=0.0009** | 0.08<br>p=0.098 | NA | Aggregate indices: Females who spent more of the time period co-residing with adult maternal sisters received more grooming. The estimate represents the expected difference in aggregate grooming indices between females that spent none of the time period with any adult daughters and females that spent 100% of the time period with an average of 1 adult maternal sister. |
| Pedigree relatedness to other adult females | 0.03<br>p=0.60 | 0.02<br>p=0.679 | 0.07<br>p=0.542 | NA | Aggregate indices: no effect. |
| Mother-daughter pair | NA | NA | NA | **1.69**<br>**p<0.0001** | Dyadic index: Mother-daughter pairs groomed more than pairs that were not mother-daughter. The estimate represents the expected difference in the dyadic grooming index for mother-daughter pairs compared to pairs who were not mother-daughter. |
| Maternal sister pair | NA | NA | NA | **0.57**<br>**p<0.0001** | Dyadic index: Maternal sister pairs groomed more than pairs that were not maternal sisters. The estimate represents the expected difference in the dyadic grooming index for maternal sister pairs compared to pairs who were not maternal sisters |
| Relatedness to partner | NA | NA | NA | **0.61**<br>**p<0.0001** | Dyadic index: Females gave more grooming to individuals they were more closely related to than individuals they were more distantly related to. The estimate represents the expected change in the dyadic grooming index as partner relatedness changed from 0 to 1. |
| **Demographic effects** |  |  |  |  |  |
| Group size | -0.01<br>p=0.124 | 0<br>p=0.309 | 0<br>p=0.72 | **-0.01**<br>**p<0.0001** | Aggregate index: No effect.<br>Dyadic index: Females gave less grooming to each social partner in larger groups. The estimates represent the expected change in each grooming index in response to an increase in group size of one group member. |
| Sex ratio | 0.05<br>p=0.646 | 0.05<br>p=0.118 | **0.09**<br>**p=0.0481** | 0.02<br>p=0.157 | Aggregate index: Females engaged in more female-female grooming interactions in groups with a more female biased sex ratio. The estimates represent the expected change in each grooming index in response to a difference between a group with an equal sex ratio versus a ration of 2 adult females to 1 adult male. |
| **Admixture effects** |  |  |  |  |  |
| Focal admixture score | 0.36<br>p=0.07 | 0.11<br>p=0.47 | 0.49<br>p=0.12 | 0.28<br>p=0.13 | Aggregate and dyadic indices: no effect |
| Partner admixture score | NA | NA | NA | 0.21<br>p=0.26 | Dyadic index: no effect |
| Focal-partner admixture score interaction | NA | NA | NA | **-0.39**<br>**p=0.01** | Dyadic index: Focals with high admixture scores groomed partners with high admixture scores more than partners with low admixture scores. Focals with low admixture scores groomed partners of all admixture scores about the same (Figure S4) |

†The top line of each row provides the effect size estimate for each fixed effect.

Including these predictors in our models allowed us to determine the effect these environmental influences had on grooming behavior, while accounting for genetic similarities between individuals in our dataset. Not only are these environmental effects interesting in their own right, but they are important to include in the animal model because if these predictors are non-randomly distributed over the pedigree, they can potentially bias the estimates of additive genetic variance for a trait. (Kruuk and Hadfield 2007; Wilson 2008).

#### Goal 2: Direct and indirect genetic effects on female grooming behavior

*Heritability of grooming given, grooming received, and total grooming using aggregate grooming indices*. For each of our three aggregate grooming indices (grooming given, grooming received, and total grooming) we used the ‘asremlr’ package in Rv.3.0.1 (Gilmour et al. 2009) to fit a series of linear mixed models with consistent fixed effect structures and increasingly complex random effect structures. We modeled the grooming behavior of individual *I* in the following series of nested models for each grooming index:

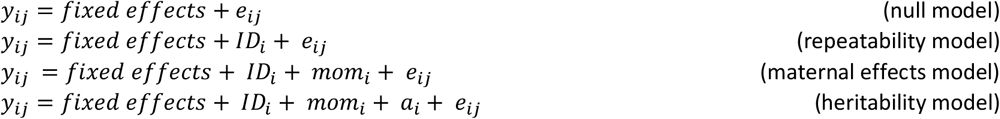

where *y*_*ij*_ is the aggregate grooming given, aggregate grooming received, or total grooming for individual *i* in year *j, e*_*ij*_ is a residual error term, *ID*_*i*_ is a random effect of the identity of the focal individual, *mom*_*i*_ is a random effect of the mother of the focal individual, and *a*_*i*_ is the additive genetic contribution of individual *i* (i.e., its breeding value). We did not include a random effect of year because the aggregate indices were standardized across years (see Supplementary Methods). We included an effect of individual identity in all models because we had repeated measures within individuals. This random effect of individual identity represents the ‘permanent environmental’ differences between individuals that arise through environmental or genetic effects. We used a likelihood ratio test to determine which of these nested models was best for each grooming index. Including the fixed effects described in Goal 1 could reduce the residual variance reported in our models which would result in increased heritability estimates. Therefore, following common practice, we report heritability estimates from models with and without fixed effects (see Results).

*Direct and indirect genetic effects on grooming given, using the dyadic grooming index*. We next fitted a series of linear mixed models using the dyadic grooming index, again using Rv3.0.1 and the ‘asremlr’ package (Gilmour et al. 2009). The primary benefit of the dyadic grooming index is that it allowed us to investigate indirect genetic effects on grooming, something that is not possible with the aggregate indices.

To determine whether indirect genetic variance contributed significantly to phenotypic variance in the dyadic grooming index, we constructed five nested models, with consistent fixed effects (as described above for the aggregate measures) and increasingly complex random effect structures. We followed the approach outlined by Wilson et al. (2011) in their investigation of indirect genetic effects for aggressive phenotypes. Specifically, we modeled the grooming given from a focal individual *i* to a grooming partner *j* in a series of five models:

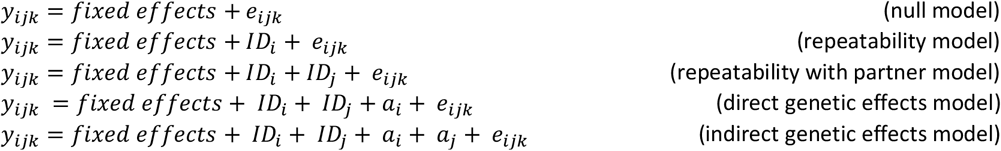

where *y*_*ijk*_ is the grooming given from individual *i* to individual *j* in year, *k*. The fixed effects, *ID*_*i*_, *a*_*i*_, and *e*_*ijk*_ terms are as described above. *ID*_*j*_ is a random effect of the individual who received grooming, and *a*_*j*_ is the additive genetic contribution of the individual who received grooming. The ‘direct genetic effects’ model allows genetic variance among the focal individuals to influence phenotypic variance in grooming given, while the ‘indirect genetic effects model’ allows genetic variance among both the focal and the partner individuals to influence phenotypic variance in grooming given by the focal partner.

As with Goal 1, because these models are nested with respect to their random effects, we used a likelihood ratio test to determine the best model for the dyadic grooming index. We tested models that included a random effects of observation year, social group, and the focal individual’s mother and found no statistically significant variation explained by these effects.

#### Goal 3: Genetic relationships between grooming given and grooming received

*Genetic covariance between grooming given and grooming received, using aggregate grooming indices*. We investigated the genetic covariance between aggregate grooming given and aggregate grooming received by using bivariate animal models. Bivariate models allow for the simultaneous estimation of sources of variance in two phenotypes as well as estimation of the sources of any covariance between these two phenotypes (see Lynch and Walsh 1998; Kruuk 2004; Wilson et al. 2010). First, to determine if the genetic covariance between aggregate grooming given and aggregate grooming received was significantly different from zero, we constructed two models, a model in which the genetic covariance was constrained to zero and a model in which the genetic covariance was free to vary (see *Example Code* in Supplement). Both models fit estimates of the environmental and maternal covariance between the traits, and the fixed effects described above. Using asremlr we compared the AIC values from these models to assess whether the genetic covariance between traits was significantly different from 0.

Next, to determine if the genetic covariance between aggregate grooming given and aggregate grooming received was significantly different from +1 (i.e., a perfect positive correlation, meaning that the two traits have the same genetic architecture), we compared the model in which the genetic covariance was free to vary to a model in which the genetic covariance was constrained to be +1. Again, both models estimated environmental and maternal covariance between the traits, and the fixed effects described above. We again compared AIC values using asremlr. When reporting our results, we rescaled the genetic covariance to a genetic correlation to facilitate interpretation of this effect.

*Covariance between DGEs and IGEs, using the dyadic grooming index*. To investigate the covariance between direct and indirect genetic effects on our dyadic index of grooming given, we fitted an additional model that estimated the covariance between these effects. To do so, we modeled a relationship between two random effects (focal breeding value and partner breeding value) so that the model fitted an unstructured 2×2 matrix, which supplies the genetic variances for the giver and receiver in the diagonal, and the covariance on the off diagonal (see *Example Code* in Supplement and McFarlane et al. 2015 for more details about this approach). To determine if this covariance was significantly different from 0 and/or significantly different from +1, we compared the AIC value from a model in which the covariance between IGEs and DGEs was free to vary to AIC values from models in which this covariance was constrained to either 0 or 1.

## RESULTS

### Goal 1: Fixed effects: Social and non-social influences on female grooming behavior

*Female grooming behavior is generally reciprocal*. The phenotypic correlation between aggregate grooming given and aggregate grooming received was statistically significant and positive, indicating that individuals who gave more grooming were likely to receive more grooming (repeated measures R=0.33, p<0.0001). The equality of grooming relationships also tended to vary systematically across an individual’s life, with females giving more grooming than they received early in adulthood and receiving more grooming than they gave at older ages. However, females showed considerable inter-individual variation in this pattern (Figure 2).

**Figure 2.**
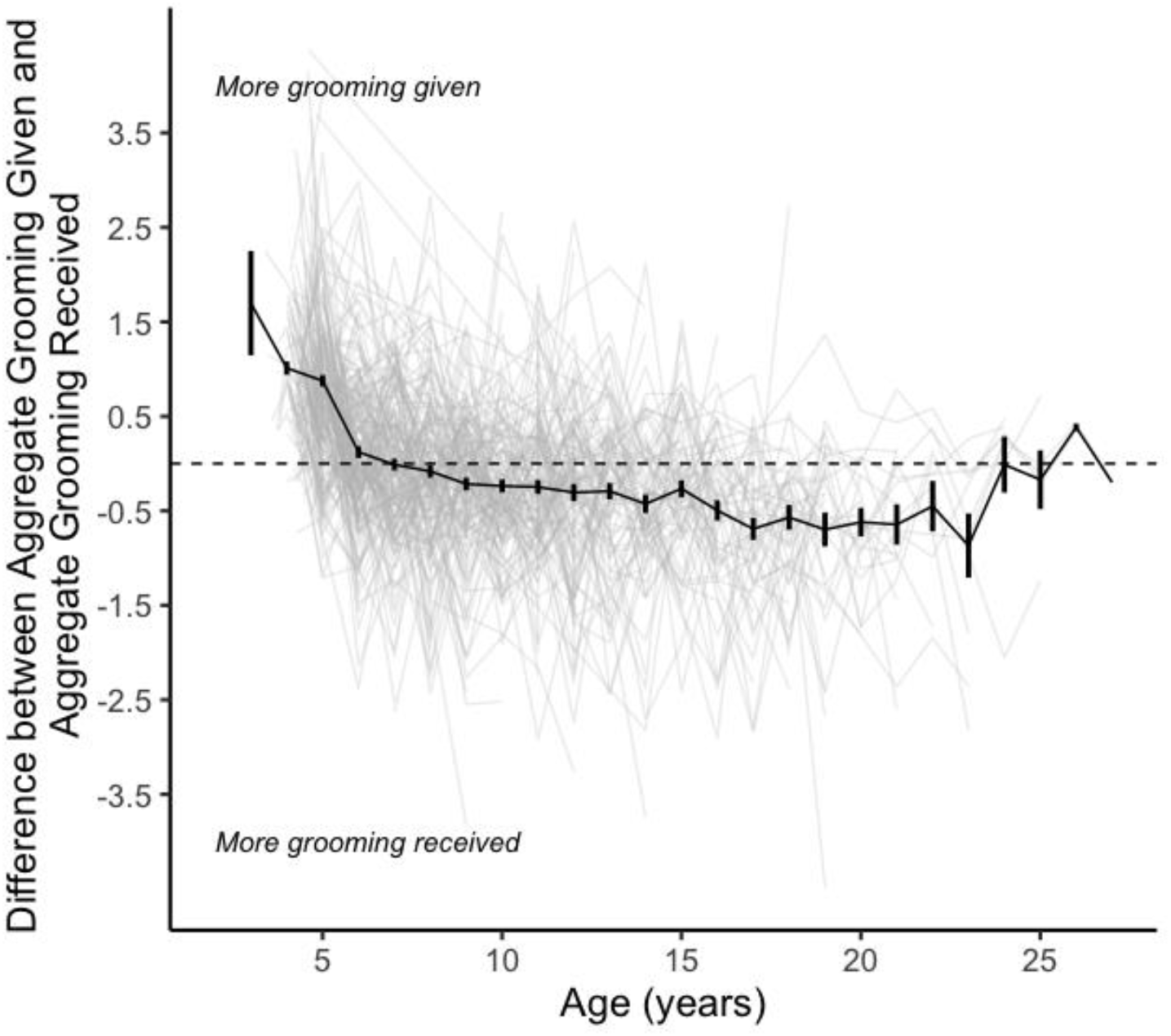
Variation in the relationship between grooming given and grooming received over the life course; each light gray line represents the values for a single female (n=224). The black line shows the average difference between grooming given and grooming received as a function of female age; bars show the standard error of the mean for each age. Values above zero represent age classes in which females gave more grooming than they received, vice versa for below zero.

*Both aggregate and dyadic grooming patterns are influenced by multiple environmental effects*. Nearly all fixed effects, apart from admixture score and pedigree relatedness to other females in the group, showed statistically significant relationships with one or more aggregate grooming indices (Table 1). Individual dominance rank was positively related to total aggregate grooming. However, dominance rank was not related to the dyadic grooming index, meaning that higher ranking individuals engaged in more total grooming activity, but not necessarily more grooming with any single partner. Grooming given and total grooming were higher for younger females and higher-ranking females, and grooming received was higher for older females. All aggregate measures of grooming (grooming given, grooming received, and total grooming) were higher for females who spent more time co-resident with their mothers and adult daughters. Females who spent more time co-resident with adult maternal sisters also received more grooming (Table 1). Group size did not influence the amount of grooming given or received, but females engaged in more total grooming interactions with other females when they were in a group with a female-biased sex ratio.

The environmental predictors of dyadic grooming given followed similar patterns as those for aggregate grooming behavior (Table 1). The effect of dominance rank on dyadic grooming involved an interaction between the dominance rank of the focal female and her partner: high-ranking females gave more grooming to high-ranking females than to low-ranking females, and low-ranking females gave more grooming to low-ranking females than to high-ranking females (Table 1, Figure S4). Females gave more grooming to their female grooming partners when they were young or when their partner was old. Individuals gave more grooming to their relatives than to non-relatives, and gave more grooming to mothers, daughters, and maternal sisters than to other partners, even when controlling for relatedness. Females gave less grooming to each female grooming partner when they were in larger groups. As with the aggregate indices, we found no simple effect of admixture score on dyadic grooming (Table 1).

However, we did detect an interaction between focal admixture score and partner admixture score: females with moderate to high admixture scores (i.e., more anubis-like females) gave significantly more grooming to other intermediate or anubis-like females than they did to yellow-like baboons (Figure S5). We did not detect a similar pattern of assortativity for more yellow-like females.

### Goal 2: Direct and indirect genetic effects on female grooming behavior

*Heritability of total grooming, grooming given, and grooming received, using aggregate grooming indices*. The heritability model was the best model for all aggregate indices of female-female grooming (Table 2, Figure 3). Our results indicate that these three phenotypes have similar heritability levels (grooming given: *h*^*2*^=0.22; grooming received: *h*^2^=0.16, total grooming: *h*^2^=0.26; Table 3). These heritability estimates represent the proportion of variance explained by additive genetic variance *after* conditioning on the fixed effects we included in our model. Conditioning on fixed effects has the potential to significantly affect heritability estimates (see Methods and Wilson 2008), therefore, we examined the role of fixed effects on our heritability estimates by running parallel models that excluded fixed effects (Table 3, final column). The slight increases in heritability when we excluded fixed effects suggest that relatives share similar values for some fixed effects, and therefore heritability is inflated without including these fixed effects in the model.

**Table 2.** Model comparisons for aggregate and dyadic grooming indices: the Heritability model was the best model (i.e., had the lowest log likelihood) for all measures of aggregate grooming, and the IGE model was the best model for dyadic grooming.

| Outcome Variable | Model | Log likelihood | p value† |
| --- | --- | --- | --- |
| **AGGREGATE INDICES** |  |  |  |
| Aggregate grooming given | Null | -673.85 | -- |
|  | Repeatability | -473.60 | <0.00001 |
|  | Maternal Effects | -471.23 | 0.029 |
|  | **Heritability** | **-464.22** | **0.0002** |
| Aggregate grooming received | Null | -680.13 | -- |
|  | Repeatability | -628.49 | <0.0001 |
|  | Maternal Effects | -623.81 | 0.0022 |
|  | **Heritability** | **-615.62** | **<0.0001** |
| Aggregate total grooming | Null | -1673.55 | -- |
|  | Repeatability | -1532.55 | <0.0001 |
|  | Maternal Effects | -1527.01 | 0.0009 |
|  | **Heritability** | **-1516.95** | **<0.0001** |
| **DYADIC INDEX** |  |  |  |
| Dyadic grooming given | Null | -7857.78 | -- |
|  | Repeatability | -7569.78 | <0.0001 |
|  | Repeatability with partner | -7497.09 | <0.0001 |
|  | Direct Genetic Effects (DGE) | -7488.44 | <0.0001 |
|  | **Indirect Genetic Effects (IGE)** | **-7482.01** | **0.0003** |

†p-values are the results of a likelihood ratio test between the given model and the model one row above

**Table 3.** Sources of variance in aggregate grooming indices from the Heritability model (see Table 2), and sources of variance in the dyadic grooming index from the Indirect Genetic Effects model (see Table 2)

| | Additive genetic contribution ($h^2$) with standard error<sup>†</sup> | Maternal contribution ($m^2$) with standard error<sup>‡</sup> | Individual permanent environment contribution ($pe^2$) with standard error<sup>§</sup> | Additive genetic contribution without fixed effects ($h^2$) with standard error |
| --- | --- | --- | --- | --- |
| **AGGREGATE INDICES** |  |  |  |  |
| Grooming given | 0.22 (0.05) | 0.06 (0.04) | 0.05 (0.06) | 0.30 (0.07) |
| Grooming received | 0.16 (0.03) | 0.007 (0.02) | 5.0e-7 (1.6e-8) | 0.18 (0.04) |
| Total Grooming | 0.26 (0.05) | 0.04 (0.03) | 2.6e-7 (1.3e-8) | 0.28 (0.07) |
| **DYADIC INDEX** |  |  |  |  |
| Focal (Direct Effects) | 0.05 (0.02) | N/A | 0.02 (0.01) | 0.03 (0.01) |
| Partner (Indirect Effects) | 0.02 (0.007) | N/A | 0.008 (0.005) | 0.01 (0.005) |

<sup>†</sup>Proportion of total phenotypic variance explained by additive genetic variance
<sup>‡</sup> Proportion of total variance explained by maternal ID variance
<sup>§</sup> Proportion of total variance explained by focal ID variance.

**Figure 3.**
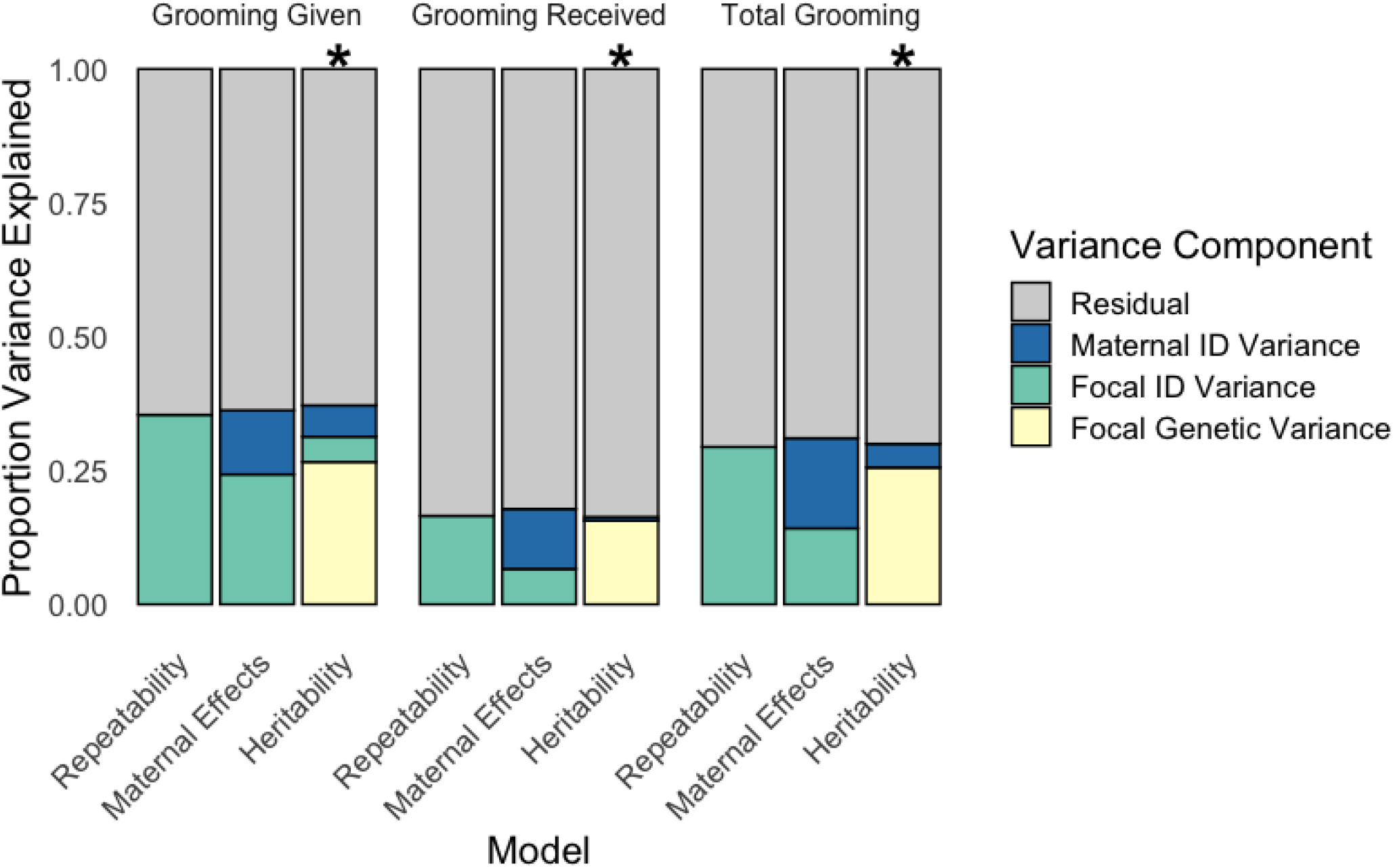
Visualization of the models of aggregate grooming depicted as the proportion of variance explained by the following random effects: direct genetic effects (additive genetic variance), the identity of a female’s mother (maternal ID variance), focal permanent environment (focal ID variance), and residual variance. For each aggregate metric, the proportion of variance explained by additive genetic variance in the “heritability” models represents the heritability. Bars with * indicate the best model as determined by a likelihood ratio test

*Direct and indirect genetic effects on grooming given, using the dyadic index*. As described above, the primary role of the dyadic index was to allow us to estimate indirect genetic effects, and indeed the IGE model was the best model among those we tested for the dyadic grooming index (Table 2, Figure 4). Because this model allowed additive genetic variance within focal *and* partner individuals to contribute to variance in grooming given, this result indicates measurable indirect genetic effects of partner identity on the amount of grooming that a focal female gave within a dyadic partnership. However, estimates of both direct and indirect genetic effects on the dyadic index were small: indirect genetic effects (i.e., genetic variation among partner individuals) explained approximately 2% of the variance in how much grooming a female gave to a particular female partner (Figure 4, Table 3). Notably, while direct genetic effects explained only 5% of the variation in the dyadic index, they explained 26% of the variance in aggregate grooming given. This large difference in the magnitude of direct genetic effects between the aggregate and dyadic models likely arises from the fact that small errors in measurement have a larger effect on our dyadic index than our aggregate index, thus increasing the residual (error) variance in our dyadic index.

**Figure 4.**
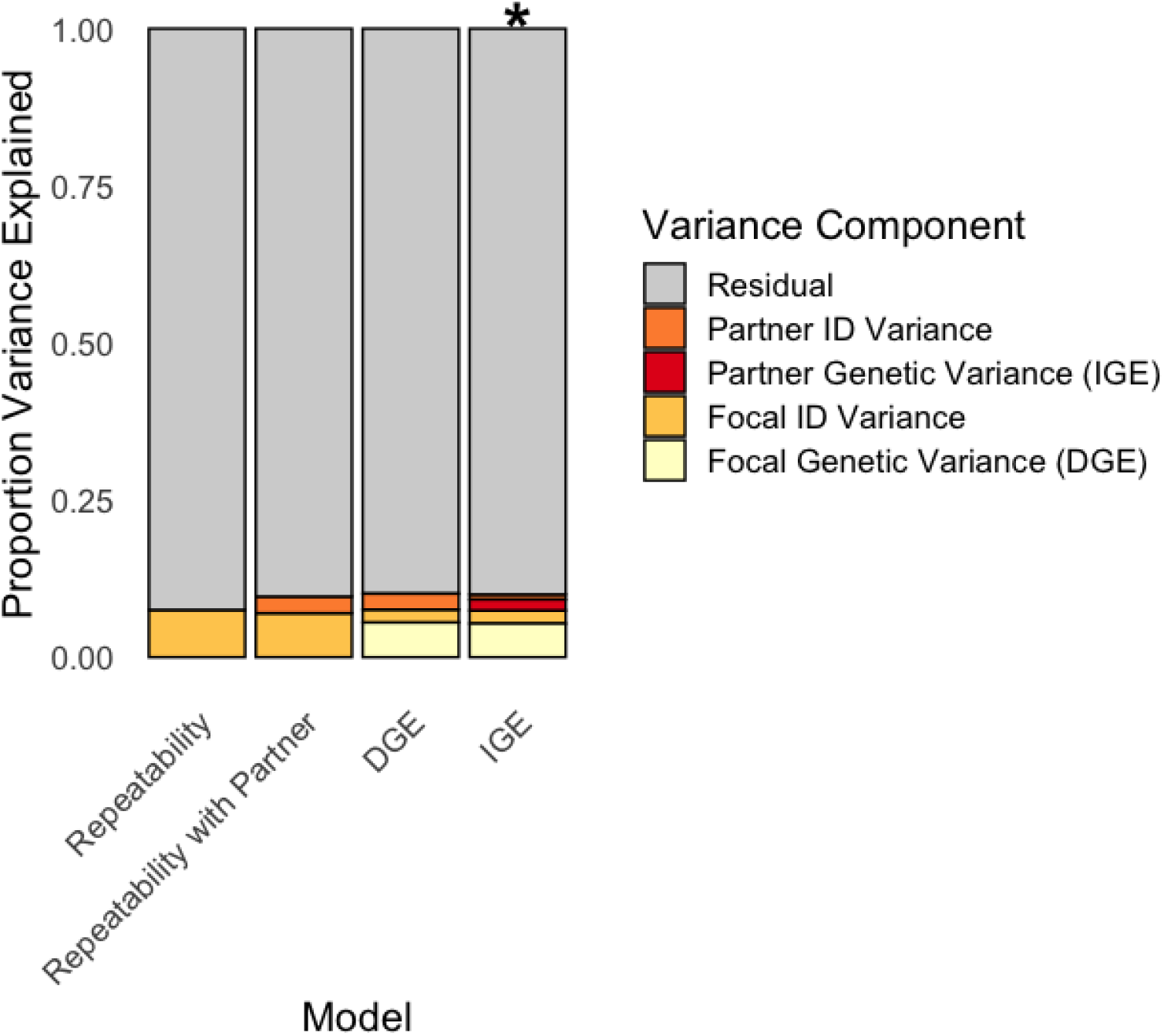
Visualization of the models of dyadic grooming depicted as the proportion of variance explained by the following random effects: direct genetic effects (focal genetic variance), indirect genetic effects (partner genetic variance), focal permanent environment effects (focal ID variance), partner permanent environment effects (partner ID variance), and residual variance. The proportion of variance explained by focal genetic variance in the DGE and IGE models represents the contribution of direct genetic effects to phenotypic variance in the dyadic metric. The proportion of variance explained by partner genetic variance in the IGE model represents the contribution of indirect genetic effects to phenotypic variance in the dyadic metric. The bar with * indicates the best model as determined by a likelihood ratio test

### Goal 3: The genetic relationship between grooming given and grooming received

*Genetic covariance between grooming given and grooming received, using aggregate grooming indices*. The genetic correlation between the phenotypes of grooming given and grooming received was strong and positive (R=0.94 ± 0.12, Table 4). Indeed, the correlation was considerably stronger than the phenotypic correlation between grooming given and grooming received (R=0.33; see above). The model with the lowest AIC value was the model with the genetic covariance between these traits fixed at 1. Our model also revealed an extremely high positive correlation (R=0.99) between the maternal effects for grooming given and grooming received (Table 4), suggesting that these two phenotypes are affected in exactly the same way by maternal effects. The model in which the genetic covariance was free to vary had a ΔAIC < 2 relative to the best model, indicating that they were statistically indistinguishable.

**Table 4.** Sources of covariance in aggregate and dyadic indices; the model with the lowest AIC is shown in the first row in each section.

| Model | Additive Genetic Correlation ($R_A$) † | Permanent Environment Correlation ($R_{pe}$)† | Maternal correlation ($R_m$)† | Residual correlation ($R_R$)† | AIC | $\Delta AIC$ |
| --- | --- | --- | --- | --- | --- | --- |
| **AGGREGATE INDICES – GROOMING GIVEN AND GROOMING RECEIVED** |  |  |  |  |  |  |
| **Genetic covariance constrained to 1** | **1** | **-0.99‡** | **0.99‡** | **0.41±0.02** | **1801.00** | --- |
| **Genetic covariance free to vary** | **0.94 ±0.12** | **-0.99‡** | **0.99‡** | **0.41±0.02** | **1802.88** | 1.88 |
| Genetic covariance constrained to zero | 0 | 0.73 ± 0.32 | 0.99† | 0.41±0.02 | 1817.49 | 16.49 |
| **DYADIC INDEX – DIRECT AND INDIRECT EFFECTS ON GROOMING GIVEN** |  |  |  |  |  |  |
| **Genetic covariance free to vary** | **0.86 ±0.06** | -- | -- | -- | **14912.27** | -- |
| Genetic covariance constrained to 1 | 1 | -- | -- | -- | 14915.83 | 3.56 |
| Genetic covariance constrained to zero | 0 | -- | -- | -- | 14975.47 | 63.2 |

†Correlations (R values) within the indices attributable to additive genetic sources (A), permanent environment sources (pe), maternal sources (m) and residual sources (R). Bolded values indicate statistically indistinguishable best models.
‡These estimates were so close to the boundaries established in model fitting (-1/1) that ASReml R was unable to provide a standard error

Interestingly, the best models (i.e., perfect or near-perfect genetic correlation between traits) also indicate a very high *negative* correlation between the effects of the permanent environment on grooming given and its effects on grooming received, while the model in which genetic covariance is constrained to zero indicates a very high *positive* correlation. Most likely, permanent environmental effects only become positively correlated when we constrain genetic covariance to zero because this model “forces” genetic effects to be fit as the consequences of the permanent environment.

*Genetic covariance between IGEs and DGEs for grooming given, using the dyadic index*. Indirect and direct genetic effects (IGEs and DGEs) for grooming given were also strongly positively correlated (R=0.86 ± 0.06, p=<0.0001). The model that allowed the genetic covariance between direct and indirect genetic effects to freely vary was the best model (i.e., had the lowest AIC, Table 4). This result is congruent with the positive genetic correlation we found between grooming given and grooming received in the aggregate models. However, the analysis of the dyadic index provides additional insight by suggesting that particular genotypes predict increased grooming in focal individuals, whether those genotypes are found in the focals themselves or in their grooming partners.

## DISCUSSION

The work presented here represents one of the first investigations into the genetic architecture of non-reproductive affiliative social behaviors in the wild. Our analysis yielded two major findings: first, the grooming behavior of adult female baboons is heritable, and second, the tendencies to give grooming and to receive grooming are strongly positively correlated, not only phenotypically but also genetically. We demonstrate the genetic correlation between the tendency to give and receive grooming in two ways: first, the amount of grooming an individual gives is genetically correlated with the amount of grooming she receives. Second, both a female’s genotype and, via indirect genetic effects, her partner’s genotype influence the relative frequency with which females groom their social partners. Specifically, the genotypes that encourage a female to give grooming *to* her social partners also appear to be genotypes that elicit grooming *from* her social partners. We found limited effects of genetic admixture on grooming behavior. We discuss our main findings below.

### Goal 1: Fixed effects: Social and non-social sources of variance in grooming behavior

The environmental and demographic factors that influence female grooming behavior have been investigated in a number of primate species, including baboons (Schino 2001; Nakamichi 2003; Lehmann et al. 2007; Akinyi et al. 2013). Our analysis is unique because, by incorporating pedigree information in the animal model, our estimates of fixed environmental effects account for pseudo-replication that may occur by including individuals with similar genetic backgrounds. Three types of environmental effects on grooming are particularly noteworthy.

*Dominance rank*. Our results are consistent with the observation, widely documented across primate species, that higher-ranking females have more grooming partners than lower-ranking females. This pattern is consistent with the well-supported hypothesis, first proposed by Seyfarth, that females groom higher ranking individuals in exchange for currencies other than grooming (e.g., agonistic support, tolerance during feeding, etc.; see Seyfarth 1977; Schino 2001). In addition, while we found no main effect of dominance rank on dyadic grooming, we did find an interaction effect, such that higher-ranking individuals gave more grooming to high-ranking partners, while lower-ranking individuals gave more grooming to low-ranking partners (Table 1). This result parallels our analysis of male-female grooming, in which the probability of grooming was highest for male-female pairs in which both partners were high-ranking (Fogel et al 2021). This result is also consistent with Seyfarth’s model, which predicts that females compete for the opportunity to groom higher ranking females, and consequently high-ranking females have the greatest access to their preferred partners (Seyfarth 1977).

*Demographic effects*. Group size did not have a significant effect on aggregate female grooming behavior, but females gave less grooming to individual partners in larger groups. Females engaged in more grooming in groups with a more female biased sex ratio. In combination, these results suggest that females in larger groups have more female grooming partners than females in smaller groups but groom each partner less when they are in a larger group, pointing towards a potential tradeoff between the strength and quantity of social bonds with females. This result is consistent with other studies that have found evidence of a decrease in group cohesion with increasing group size (Dunbar 1991; Henzi et al. 1997; Lehmann et al. 2007; Cheney et al. 2012).

*Effects of individual admixture on grooming*. Our measure of genetic admixture had no statistically significant effect on any measure of aggregate grooming, although we detected a trend towards more admixed individuals giving more grooming (Table 1). However, in the dyadic grooming index, we detected a significant interaction between the admixture score of the focal female and her partner: individuals with more anubis ancestry preferentially groomed partners that were more anubis-like, while more yellow-like females appeared insensitive to the ancestry of their partners. Similar effects of admixture score have been described for male mating success and for male-female affiliative behavior in our study population (Tung et al. 2012, Fogel et al. 2021), together suggesting that anubis-like females may behave more assortatively (with respect to ancestry) than yellow-like females in their choice of social or mating partners. This phenomenon could arise either through competitive access to preferred partners or because of differences in preference strength (e.g., if the costs of interacting with a member of a different sub-population are higher for more anubis-like females than for yellow-like females). Alternatively, as the most anubis-like animals in our population tend to be products of admixture in the past few generations, they may express an assortative preference that has been eliminated in animals that are the descendants of much older admixture (e.g., if such selectivity bears a cost).

*Positive correlation between aggregate grooming given and received*. Aggregate grooming given and aggregate grooming received are positively phenotypically correlated in the Amboseli baboons (repeated measures R=0.33, p<0.0001). This correlation is likely driven by positive genetic covariance and positive maternal effect covariance (Table 4), as our best models identify strong correlations in both cases, combined with a negative correlation for permanent environmental effects.

### Goal 2: Direct and indirect genetic effects on female grooming behavior

We found that the tendency to engage in affiliative social interactions with other females is heritable and consequently, may evolve in response to natural selection. This provides an important conceptual link between studies that have demonstrated apparent fitness benefits of social interactions, and studies that have demonstrated heritability for phenotypes that influence social interactions (e.g., physiology: Insel and Shapiro 1992; Walum et al. 2008; Staes et al. 2018; personality: Jang et al. 1996; Brent et al. 2014; Staes et al. 2016; morphology: Moore 1990; Schielzeth et al. 2012). However, further work is needed to predict the magnitude and direction of this response. While we have strong evidence linking grooming behavior to both health (Akinyi et al. 2013) and survival in this study population (Silk et al. 2003; Archie et al. 2014; Campos et al. 2020), we do not yet know whether grooming behavior has a causal link to survival, or is simply correlated with other traits that do.

The highest heritability we detected was 0.26 for total grooming, consistent with heritability estimates reported for life history and behavioral traits in wild populations, but lower than generally reported for morphological traits (Visscher et al. 2008, Houslay et al. 2021). This value may indicate that the quantitative genetics of grooming is more like that of life history traits than morphological traits, and that selection has effectively reduced genetic variation for grooming behavior, resulting in a relatively small numerator (V_A_) in the calculation of heritability. Alternatively, grooming may be responsive to many environmental inputs and as a result may have a relatively large phenotypic variance, V_P_ – the denominator in the calculation of heritability (Mousseau and Roff 1987). The well-established association between grooming behavior and survival, as well as the complex nature of behavioral phenotypes suggest that both processes could be involved in the heritability of grooming behavior.

Our estimates of indirect genetic effects were small but measurable, accounting for 2% of the variance in how much grooming a female gave to a particular female partner. In spite of the small magnitude of this effect, our results are the first demonstration, to our knowledge, of indirect genetic effects on affiliative social behaviors in a wild vertebrate population. IGEs are thought to be of particular importance in the evolution of social behavior compared to other phenotypes (Wolf et al. 1998; Moore et al. 2002; Cheverud 2003; McGlothlin et al. 2010, Bailey et. al 2018), and our study stands as an important example of the feasibility of measuring IGEs for social behavior in the wild.

### Goal 3: The genetic relationship between grooming given and grooming received

What explains the very strong genetic correlation between grooming given and grooming received (R=0.94 ± 0.12, Table 4)? Grooming is near-certain to be a highly polygenic trait, similar to the majority of other complex traits that have been investigated in the post-genomic era, so simple linkage disequilibrium is unlikely to account for this magnitude of covariance (Flint 2003, Boyle et al. 2017). We propose two alternative explanations. First, the high genetic correlation we observe may result from the fact that female baboons both give grooming to and receive grooming from close relatives more often than they do with distant or non-relatives (Silk et al. 2006b,a). While we accounted for this familial effect by modeling the presence of the focal’s maternal kin and her overall relatedness to her social group, this approach may fail to fully account for how the presence of close relatives increases a female’s grooming rates, both given and received. If so, our estimate of the genetic covariance between these traits may be inflated. For instance, the tendency for relatives to preferentially groom each other will mean that an individual’s phenotype for grooming given is a stronger predictor of her relatives’ phenotypes for grooming received than of the phenotypes of her non-relatives (and vice versa). Under such a scenario, the phenotypic correlation between relatives would be attributed to genetic covariance between grooming given and grooming received. Indeed, when family structure influences a phenotype, it can be extremely difficult (without cross-fostering) to fully disentangle the environmental effects of relatedness from the genetic ones (Kruuk and Hadfield 2007).

A second alternative explanation for the strong genetic correlation between grooming given and received is that these two phenotypes may not truly be separate, but may instead emerge from the same underlying, partially heritable trait. One candidate trait would be the tendency to reciprocate when groomed. As noted earlier, individuals tend to form highly reciprocal grooming relationship in many primate species (Schino and Aureli 2008), and in baboons the most enduring social relationships are the most reciprocal ones (Silk et al. 2006a, 2010). It is possible that our grooming data do not reflect the tendency to give and receive grooming *per se*, but instead reflect the tendency to reciprocate when groomed. That is, given that individual A begins a grooming relationship with individual B at some point in its life, it is possible that much of the grooming we subsequently measure between A and B depends on each individual’s tendency to reciprocate grooming. If individuals assort socially according to their tendency to reciprocate (so that high reciprocators tend to prefer each other), the result would be a very strong positive genetic correlation between grooming and being groomed. This interpretation is supported by the extremely high correlation between the maternal effects for grooming given and grooming received, indicating that the two phenotypes are affected in exactly the same way by maternal effects, supporting the idea that both emerge from a single trait (Table 4). This interpretation is also supported by the positive genetic correlation between direct and indirect effects on grooming given (Table 4), which indicates that a given genotype will give and elicit similar amounts of grooming from/to its grooming partners.

An additional strategy for investigating whether the genetic correlation between grooming given and grooming received can be explained by reciprocity would involve trait-based investigations of indirect genetic effects (see Wolf et al. 1998; Bleakley and Brodie IV 2009; McGlothlin and Brodie 2009). Trait-based approaches focus on how phenotypes are influenced by specific traits in a social partner, as opposed to simply estimating the proportion of variance in the focal phenotype explained by similarity in the partner’s genotype, as we did here (see also Wilson et al. 2005, 2009, 2011; Sartori and Mantovani 2012). Our approach, a ‘variance-partitioning method,’ is useful for initial estimates of direct and indirect genetic effects and genetic covariance, and is well suited to the genetic structure of our natural breeding population. Future analyses using a trait-based approach would generate further insight into the mechanistic basis of the observed reciprocity. However, it requires fine-grained phenotypic data on the duration and sequential order of grooming bouts, which is not a part of our standard behavioral data collection protocol.

If the strong genetic covariance between grooming given and grooming received does indeed reflect a close genetic linkage or even a genetic identity between these two apparently distinct traits, our results would suggest a role for genetic architecture in the evolution of cooperation and reciprocity in primates. This phenomenon has been documented in microbes such as the social amoeba *Dictyostelium discoideum* (see also, Rainey and Rainey 2003; Griffin et al. 2004; Xavier and Foster 2007; Springer et al. 2011). Under certain conditions, some *D. discoideum* cells die to form a stalk that facilitates the dispersal of other cells in reproductive spores (Strassmann et al. 2000). This pattern of stalk formation is often interpreted as an act of extreme cooperation and even altruistic sacrifice. Genetic architecture, namely pleiotropy, has been implicated in preventing cheaters who avoid the sacrifice of stalk formation from achieving the reproductive benefits of spore production. Foster et. al (2004) showed that the *dimA* gene is required for both differentiation into the cooperative stalk, and for correct allocation to the reproductive spore. The pleiotropic effects of this gene mean that cheating genotypes that avoid the sacrifice of the cooperative stalk also fail to allocate correctly to the reproductive spore. This genetic architecture serves to facilitate the evolution of cooperation by preventing the spread of cheaters.

The strikingly high genetic covariance between grooming given and grooming received we report here suggests that mechanisms similar to those described for *Dictyostelium discoideum* could potentially be at work in multicellular social organisms. Specifically, strong genetic linkages between reciprocity-related phenotypes may make it difficult for cheaters (e.g., those who do not give grooming in response to receiving it) to emerge and invade. In this scenario, the strong genetic correlation between the tendency to provide grooming and the tendency to elicit grooming from social partners would have an effect similar, in principle, to the pleiotropic *dimA* effect in *D. discoideum*.

### Future directions

The work described here represents an important step forward in the integration of primate behavioral ecology and quantitative genetics. We hope this integration serves to advance both fields, as behavioral ecology investigates how behavior might evolve in response to ecological and environmental pressures, and quantitative genetics provides the information needed to build realistic evolutionary models that consider the genetic (co)variation in traits (Cheverud and Moore 1994).

This work is also a rare example of an analysis of both genetic variance and indirect genetic effects in affiliative social behavior in a wild vertebrate. As such, it yields several prospects for further investigation. First, the measurable heritability of grooming behavior—a trait previously linked to survival—motivates a more detailed analysis of the magnitude of the phenotypic response to selection on grooming behavior. The theoretical potential for IGEs and for IGE-DGE covariance to fundamentally shape the evolution of social traits has been well documented (e.g., Wilson et al. 2009, Bijma and Wade 2008, McGlothlin al 2010), but few empirical studies have estimated the complete set of necessary parameters to predict how social traits respond to selection. These parameters include DGEs, IGEs, their covariance, as well as relatedness within the group, group size and measures of direct and social (or individual and group level) selection gradients (Bijma and Wade 2008). We have laid the groundwork for such an investigation here by estimating the relevant quantitative genetic parameters. Estimates of relevant selection gradients are still needed for understanding short-term evolutionary dynamics of grooming; these will become increasingly feasible as data collection at this long-term field study continues.

Second, among our environmental effects, the tendency for more anubis-like females to groom more anubis-like partners is intriguing, especially given that admixture scores have no detectable effect on aggregate grooming patterns. This result contributes to growing evidence of subtle behavioral differentiation between yellow and anubis baboons (Tung et al. 2012, Franz et al. 2015, Fogel et al. 2021), which are generally viewed as behaviorally and ecologically similar. Finally, the strong genetic correlation between grooming given and grooming received demands further investigation to determine whether this correlation reflects the genetic architecture of this trait, or the fact that family effects on grooming phenotypes are not fully accounted for in our models. Either result will have interesting implications; the former because of the light it sheds on the evolution of affiliative grooming, and the latter because of the methodological challenge it raises for us and other researchers who study affiliative behavior that occurs among both kin and non-kin.

## Acknowledgments

We gratefully acknowledge the support of the National Science Foundation and the National Institutes of Health for the majority of the data represented here, currently through NSF IOS 1456832, NIH R01AG053308, R01AG053330, R01AG071684, R01HD088558, R01AG075914, and P01AG031719. E.M.M was supported by NSF IOS 1501971. We thank Duke University, Princeton University and the University of Notre Dame for financial and logistical support. In Kenya, our research was approved by the Kenya Wildlife Service (KWS), the Wildlife Research & Training Institute, the National Environment Management Authority (NEMA), and the National Council for Science, Technology, and Innovation (NACOSTI). We also thank the University of Nairobi, the Institute of Primate Research, the National Museums of Kenya, the members of the Amboseli-Longido pastoralist communities, the Enduimet Wildlife Management Area, Ker & Downey Safaris, Air Kenya, and Safarilink for their cooperation and assistance in the field. Particular thanks go to the Amboseli Baboon Research Project field team (R.S. Mututua, S. Sayialel, J.K. Warutere, I.L. Siodi, G. Marinka, B. Oyath) and camp staff, to T. Wango and V. Oudu for their untiring assistance in Nairobi, and to Jeanne Altmann for her fundamental contributions to the Amboseli baboon research. The baboon project database, BABASE, was designed and programmed by K. Pinc and is expertly managed by N.H. Learn and J.B. Gordon. For a complete set of acknowledgments of funding sources, logistical assistance, and data collection and management, please visit http://amboselibaboons.nd.edu/acknowledgements/.

## Notes

### Competing Interest Statement

The authors have declared no competing interest.

