## Supplemental Material for "Genetic variance and indirect genetic effects for affiliative social behavior in a wild primate"

**Supplementary Methods**

*Study population and grooming data collection*

Both yellow and anubis baboon populations are subdivided into stable social groups, which each contain multiple adults and juveniles of both sexes, ranging in size from approximately 20-100 animals. The Amboseli Baboon Research Project monitors multiple such groups (‘study groups’) in the Amboseli ecosystem of Kenya. Four of our study subjects were born before continuous observation of our study population began and were first observed as juveniles or young adult females. Their ages were estimated based on patterns of growth and development (Altmann et al. 1981). Ages of all other subjects were known to within a few days’ error (n=204) or to within a few months’ error (n=16). We excluded a few time periods during which rates of behavioral data collection were low because of logistical challenges in the field. We also excluded periods of fission/fusion between social groups. For individual females and dyads, we excluded years during which the female (for aggregate grooming measures) or the dyad (for dyadic grooming measures) was present in the study population for less than 60 days; these included years in which a female reached maturity or died partway through the year.

Our grooming data were collected on all observed grooming events between adult females within the group. However, some grooming events were likely missed, especially in large groups. Uneven sampling of individuals was avoided by collecting the great majority of our grooming data during the course of random-order focal animal sampling on adult females and juveniles. The collection of grooming data was not restricted to grooming by the focal animal because the randomization of observer effort with respect to focal females ensured that observers continually moved to new locations within the group and observed all adult females and juveniles on a regular rotating basis.

*Relationship between grooming frequency and observer effort*

From our observations of grooming behavior, we calculated the daily frequency with which each study subject female was observed giving grooming to any other adult female in the population and the daily frequency with which she was observed receiving grooming from any other adult female in the population. Because observer effort (the number of person-hours we devote to observations of each group) does not increase with increasing group size, the apparent rate of both grooming given and grooming received is higher in smaller social groups than in larger groups. This difference arises as a simple artifact of having a constant number of observers regardless of group size. To correct for this artifact, we regressed daily rates of grooming (given or received) for all adult females alive in the population during the same year against observer effort, measured as the number of focal animal samples collected in a group per adult female per day. Specifically, we conduct 10-minute focal animal samples during daily observations, rotating through adult females in random order, completing each rotation before we begin another. Thus, the number of focal animal samples per female per day changes systematically as a function of group size in a manner that reflects the amount of time we spend directly observing each female, and is a good measure of per-female observer effort (see Archie et. al 2014 for details)

Using the residuals of grooming frequency on observer effort to define our phenotypes of interest imposes some limitations on our analysis, but also provides distinct benefits. The primary limitation of using residuals in analyses is that the parameter estimates for any fixed effects that are truly correlated with observer effort will be conservatively biased (Darlington and Smulders 2001). In our case, these fixed effects include group size and ordinal dominance rank. Despite this limitation, we chose to use residuals as our phenotype of interest instead of raw grooming counts because this approach makes our results easier to interpret in two important ways. First, the data are distributed in an intuitive way: positive values indicate more grooming behavior than the population mean for that time period while negative values indicate less grooming behavior than the population mean. Second, the quantitative genetic parameter estimates produced by a linear mixed effects model are significantly easier to interpret than the parameters produced by a generalized linear mixed effects model (which we would need to implement if our response variable was uncorrected counts of grooming bouts) because GLMMs provide inference on a statistically convenient latent scale. However, we wish to express quantitative genetic parameters on the scale upon which traits our traits were measured. While methods exist for converting parameters expressed on the latent scale to the observed scale (see de Villemereuil et al. 2016) they are not without complication and because our primary interest is in understanding genetic contributions to grooming behavior, and not in the effect of group size on grooming behavior, we chose the statistical approach that provides the most easily interpreted quantitative genetic parameters.

*Aggregate measures of grooming*

Aggregate index of total grooming

Following Archie et. al 2014, we used an aggregate index of total grooming as our measure of social integration (this is the same metric as the female social connectedness index in Archie et. al 2014). To create this index, we first calculated the annual mean value of the female’s residuals from a regression of daily rates of grooming given on observer effort (see above) and the annual mean value of the female’s residuals from a regression of daily rates of grooming received on observer effort (see above). These two annual mean values were based not on the calendar year, but on each female’s birth date and hence age class (e.g., her 6^th^ year of age, 7^th^ year of age, and so on); this approach allows us to compare age-specific grooming behaviors. For each female, the yearly aggregate index of total grooming was the average of her mean value for grooming given and her mean value for grooming received.

Aggregate indices of grooming given and grooming received

The aggregate indices of grooming given and grooming received were simply the component metrics used to calculate the aggregate index of total grooming, described above.

*Dyadic measure of grooming*

To explicitly investigate the role of indirect genetic effects on grooming behavior, we calculated a dyadic grooming index for each pair of adult females that were co-resident in a social group for at least 60 days during the study period (January 1983-June 2017) and that had at least one grooming interaction. We used the same set of females and female years as in the aggregate indices. However, for ease of calculation, we calculated the dyadic index for each calendar year instead of for each year of age for a given female. The dyadic grooming index was based on the daily rate of grooming given by the focal individual to a specific partner over the course of a calendar year, considering only the days where both focal and partner were present as adults in the same social group, and thereby available to each other as grooming partners. As with the aggregate indices described above, the dyadic index is the residual of the regression of this daily grooming rate on observer effort. Positive values indicate dyads in which the focal gave above average amounts of grooming (relative to the mean of the population, controlling for observer effort) and negative values indicate dyads where the focal gave less grooming than average (controlling for observer effort) as compared to the entire population. Hence, dyadic grooming indices do not have to be symmetrical between the two females in a dyad. In our mixed effects models, we included only dyads in which both members of the dyad were in our set of study subjects.

*Phenotypic correlation between aggregate grooming given and aggregate grooming received*

We used the ‘rmcorr’ package in R v3.4.1 to calculate the phenotypic correlation between our repeated measures of aggregate grooming given and aggregate grooming received. Rmcorr uses analysis of covariance to statistically adjust for inter-individual variability and estimate the linear fit between aggregate grooming given and aggregate grooming received for each individual, accounting for repeated measures on individuals (Bland and Altman 1995a, 1995b).

*Appropriateness of the animal model*

Pedigree structure

In this population, maternities are identified from long-term records of births, and maternities and paternities are verified with genetic parentage analysis, using microsatellite genotypes obtained from DNA derived from fecal samples or, in some cases, blood samples. Specifically, analyses of paternity and relatedness are routinely conducted for the study population (Buchan et al. 2003, Alberts et al. 2006, Charpentier et. al 2008, Tung et. al 2012). For samples extracted from faeces, all apparent homozygous genotypes are reamplified at least four and up to seven additional times to guard against allelic dropout. All genotype data were produced on either an ABI 3700 Sequence Analyzer or an ABI 3730xl Sequence Analyzer.

The maternities of all our study subjects are known, but only 77% of the paternities are known. While the pedigree for our entire study population consists of over 1500 individuals, the subset of this pedigree that describes all known relationships between all 224 of our study subjects onsists of only 347 individuals. This smaller pedigree has 209 identified father-offspring relationships, 274 identified mother-offspring relationships, and a maximum of 6 generations within a family. It includes 225 maternal half-sibling pairs, 320 identified paternal half-sibling pairs (though some paternal siblings in our dataset may be undetected) and 20 full sibling pairs. The average relatedness between any two individuals in our trimmed pedigree is 0.014, although this may be an underestimate, given some missing paternal links.

Grooming patterns among kin

Most female-female grooming in baboons occurs between maternal relatives (Silk 1987). Interactions between relatives can present challenges for the animal model, particularly in partitioning between direct and indirect sources of genetic variance. Specifically, if interactions occur only between individuals who are equally related (e.g., when interactions occur exclusively within sib-families), the direct and indirect genetic variance are not statistically distinguishable (see Bijma 2014, appendix in Bijma et al. 2007, Cheng et al. 2009). However, this problem does not affect our dataset. Female baboons have strong grooming bonds with close relatives, but they groom with individuals of all levels of relatedness (Silk et al. 2006*a*,*b*, Fig S2). Most of the grooming pairs in our study are not closely related, though more grooming events occur per pair in the more closely related dyads (see results in the main text). In our sample, relatedness between grooming pairs ranges from 0 to >0.5 (Figure S1), and a given pair of half siblings often has only partially overlapping sets of grooming partners (Figure S3). Indeed, because of the multi-male, multi-female mating system of baboons, even when maternal siblings share the same grooming partners, they often have different pedigree relationships with their grooming partners (Figure S3). This combination of differential interactions and relatedness with social partners among maternal half siblings allows the animal model to partition between direct and indirect genetic effects.

Because female-female grooming is most common between maternal relatives, the presence or absence of these relatives is an important predictor of female grooming. In our aggregate grooming models, we include the number and type of relatives present as fixed effects (see below and Supplementary Methods for details) and in our dyadic grooming models we include the pedigree relatedness and type of relationship (e.g., mother-daughter pair) as fixed effects.

Accounting for genetic admixture

As noted in the main text, the study population is composed of hybrid individuals who tend to have majority yellow baboon ancestry but also carry some introgressed ancestry from neighboring anubis baboon populations (Alberts and Altmann 2001; Tung et al. 2008). Recent work suggests that intermittent gene flow has been occurring between our study population and neighboring anubis populations for hundreds to thousands of generations (Wall et. al, 2016). Using the same 7-14 microsatellite loci that we used to construct the pedigree, we calculated the proportion of recent anubis versus yellow ancestry in each of our study subjects using Structure 2.3 (Pritchard et al. 2000, Falush et al. 2003; see Tung et al. 2008; Charpentier et al. 2012 for details of its use in the baboon study). Because admixed individuals are fully viable and reproduce freely in Amboseli the amount of recent admixture varies continuously across individuals in our population from yellow-like to anubis-like (Vilgalys et al. 2022); here we used microsatellite-based scores because resequencing data are not available for many individuals in our sample). In our study subjects, the mean point estimate for the microsatellite-based admixture scores was 0.25 (range 0.024 to 0.899). We include admixture score as a fixed effect in order to capture genetic variance in grooming behavior that is explained by anubis versus yellow ancestry. Consequently, our estimates of heritability from models that include fixed effects are an estimate of the proportion of phenotypic variance in our population that is due to genetic variance *independent of* admixture between yellow and anubis genetic backgrounds.

*Example Code*

Bivariate models for aggregate indices

To determine the genetic correlation between grooming given and grooming received, we constructed three models: a model in which the genetic covariance was constrained to zero, a model in which the genetic covariance was constrained to +1 and a model in which the genetic covariance was free to vary. The code for these models will be difficult to interpret without some familiarity with Asreml-R. Here, we have highlighted the primary differences between these three models, each presented below, with red text. Please see the [Asreml-R manual](https://www.vsni.co.uk/downloads/asreml/release2/doc/asreml-R.pdf) for details about the rest of the code. This code is for Asreml-R version 4.

#### Cartoon model with genetic covariance constrained to zero:

```
asreml(fixed=cbind(trait1,trait2)~trait+trait:fixed_effects,
       random=~corgh(trait, init=c(-0.01,.11,.11)):sname
              +corgh(trait, init=c(0.11,0.11,.11)):vm(ANIMAL, ainv*)
              +corgh(trait, init=c(.08,.11,.11)):mom,
       residual=~units:corgh(trait, init=c(0.1,0.1,0.1)),
       data=data,
       G.param=init.zero**
)
```

*ainv is determined from the pedigree file

**init.zero is a set of parameters with the covariance between genetic variance for actor and genetic variance for actee set to zero

#### Cartoon model with genetic covariance constrained to positive 1:

```
asreml(fixed=cbind(trait1,trait2)~trait+trait:fixed_effects,
       random=~corgh(trait, init=c(-0.01,.11,.11)):sname
              +corgh(trait, init=c(0.11,0.11,.11)):vm(ANIMAL, ainv)
              +corgh(trait, init=c(.08,.11,.11)):mom,
       residual=~units:corgh(trait, init=c(0.1,0.1,0.1)),
       data=data,
	 G.param=init.one***
)
```

***init.one is a set of parameters with the covariance between genetic variance for actor and genetic variance for actee set to zero

#### Cartoon model with genetic covariance free to vary:

```
asreml(fixed=cbind(trait1,trait2)~trait+trait:fixed_effects,
       random=~corgh(trait, init=c(-0.01,.11,.11)):sname
              +corgh(trait, init=c(0.11,0.11,.11)):vm(ANIMAL, ainv)
              +corgh(trait, init=c(.08,.11,.11)):mom,
       residual=~units:corgh(trait, init=c(0.1,0.1,0.1)),
       data=data
)
```

The primary difference in these models is no assignment of fixed values for G.param in the model where the covariance is free to vary.

Covariance between direct and indirect genetic effects with the dyadic index

To determine the genetic correlation between the direct genetic effects of grooming given to a partner and the indirect genetic effects of grooming elicited from a partner, we fitted three models: a model in which the genetic covariance was constrained to zero, a model in which the genetic covariance was constrained to +1 and a model in which the genetic covariance was free to vary. The code for these models will be difficult to interpret without some familiarity with Asreml-R. Here we have highlighted the primary differences between the models with red text, please see the [Asreml-R manual](https://www.vsni.co.uk/downloads/asreml/release2/doc/asreml-R.pdf) for details about the rest of the code. This code is for Asreml-R version 4.

#### Cartoon model with the DGE/IGE covariance constrained to zero:

```
asreml(fixed=trait1~fixed_effects,
random=~str(~vm(actor,ainv*)+vm(actee,ainv),~corgh(2):vm(actor,ainv))
+ide(actor)+ide(actee),
      data=data,
      G.param = initial_values.zero**
)
```

*ainv is determined from the pedigree file

**initial_values.zero is a set of parameters with the covariance between genetic variance for actor and genetic variance for actee set to zero

#### Cartoon model with the DGE/IGE covariance constrained to positive one:

```
asreml(fixed=trait1~fixed_effects,
random=~str(~vm(actor,ainv*)+vm(actee,ainv),~corgh(2):vm(actor,ainv))
+ide(actor)+ide(actee),
      data=data,
      G.param = initial_values.one***
)
```

*** initial_values.zero is a set of parameters with the covariance between genetic variance for actor and genetic variance for actee set to one

#### Cartoon model with DGE/IGE covariance free to vary:

```
asreml(fixed=trait1~fixed_effects,
random=~str(~vm(actor,ainv*)+vm(actee,ainv),~corgh(2):vm(actor,ainv))
+ide(actor)+ide(actee),
      data=data,
)
```

The primary difference in these models is no assignment of fixed values for G.param in the model where the covariance is free to vary.

|  | **Aggregate Grooming Indices** | **Dyadic Grooming Index** | **Variable Type** |
| --- | --- | --- | --- |
| **Individual traits** |  |  |  |
| Focal individual’s ordinal dominance rank^a^ | Included | Included | Continuous |
| Partner individual’s ordinal dominance rank^a^ | NA | Included | Continuous |
| Focal-partner rank interaction | NA | Included | Interaction |
| Focal age at start of observation period^b^ | Included | Included | Continuous |
| Partner age at start of observation period^b^ | NA | Included | Continuous |
| Focal-partner age interaction | NA | Included | Interaction |
| **Family effects** |  |  |  |
| Proportion of time period co-resident with mother^c^ | Included | NA | Continuous |
| Proportion of time period co-resident with adult daughters^d^ | Included | NA | Continuous |
| Proportion of time period co-resident with adult maternal sisters^e^ | Included | NA | Continuous |
| Total pedigree relatedness to other adult females^f^ | Included | NA | Continuous |
| Mother-daughter pair | NA | Included | Categorical (yes/no) |
| Maternal sister pair | NA | Included | Categorical (yes/no) |
| Relatedness to partner^g^ | NA | Included | Continuous |
| **Demographic effects** |  |  |  |
| Group size^h^ | Included | Included | Continuous |
| Sex ratio^i^ | Included | Included | Continuous |
| **Admixture effects** |  |  |  |
| Focal admixture score^j^ | Included | Included | Continuous |
| Partner admixture score^j^ | NA | Included | Continuous |
| Focal-partner admixture score interaction | NA | Included | Interaction |

**Table S1. Fixed effects used in models.**

^a^Social dominance rank is calculated on a monthly basis by minimizing entries below the diagonal in agonism matrices (see Lea et al. 2014).

^b^Female age can be determined with a high degree of certainty, because in most cases we know female birthdates to within just a few days’ error.

^c^The proportion of the year that an individual was co-resident with her mother. Co-residency means the mother and daughter were alive and in the same social group

^d,e^The proportion of the year that an individual was co-resident with her adult daughter^e^ or maternal sisters^e^. If an individual was co-resident with more than one adult daughter or adult maternal sister we summed the percentage of time spent with each daughter or sister, meaning these values can exceed 1.

^f^The sum of the simple pedigree relatedness values between the focal individual and all other adult females in the group (except for her mother, adult daughters and adult maternal sisters, who were modeled separately). This sum was weighted by how many days she spent with each individual in the given time period.

^g^Pedigree relatedness between the dyad

^h^The average number of adults (females who have reached menarche plus males who have achieved adult dominance rank) in the social group over the year

^i^The average number of adult females present over the course of the year divided by the average number of adult males present over the course of the year. Higher values indicated time periods where the group was female-biased.

^j^As noted in the main text, the study population has majority yellow ancestry, with some contribution from anubis baboons (Alberts and Altmann 2001; Tung et al. 2008). We have calculated admixture scores based on microsatellite data, where higher scores represent more anubis-like ancestry (see Tung et al. 2008, Tung et al. 2012 for details).

**Supplementary Figures**

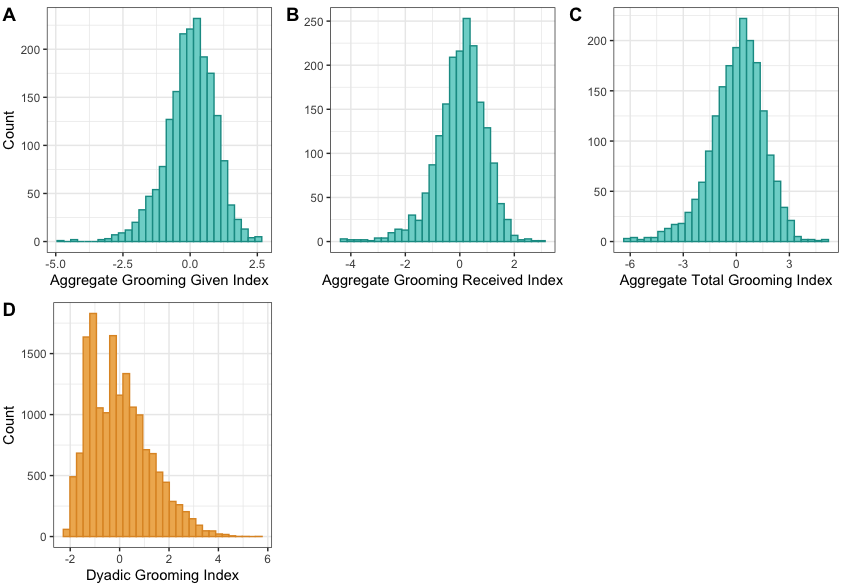

**Figure S1:** Distributions of grooming index values. (A) The index of *aggregate grooming given* reflects the frequency with which a given female in a given year of life groomed other adult females, relative to the grooming given by all other adult females alive in the same year. (B) The index of *aggregate grooming received* reflects the relative frequency with which a given female in a given year received grooming from other adult females. (C) The *aggregate index of total grooming* is the average of the *aggregate grooming given* and *aggregate grooming received* indices, calculated for each adult female for each year of her adult life. Positive values for any of these indices indicate females with relatively high frequencies of grooming (given, received, or aggregate) for the population in that year; negative values represent females with relatively low frequencies of grooming for that year. (D) The dyadic grooming index was calculated for each adult female dyad for each calendar year in which both the focal and partner were present as adults in the same social group and thereby available to each other as grooming partners. Positive values of the dyadic index indicate cases in which an adult female gave high frequencies of grooming to a specific partner relative to all other partner pairs in the population for that year, while negative values indicate cases in which an adult female gave relatively low frequencies of grooming to a specific partner.

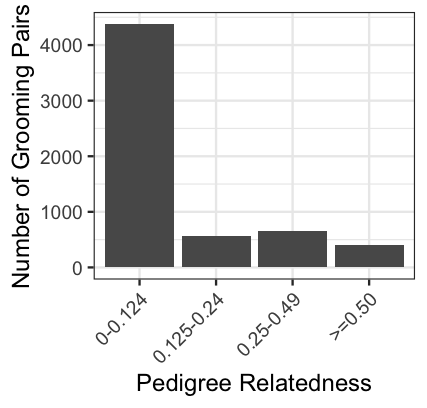

Figure S2 Distribution of grooming pairs across relatedness categories. Because our pedigree is up to six generations deep and we have both maternal and paternal connections, some pairs of individuals fall in between the typical categories of pedigree relatedness. For instance, we have 544 pairs with r=0.25 and 397 pairs with r=0.5, and we also have 105 pairs for which r falls between 0.25 and 0.5, and 6 pairs for which r>0.5.

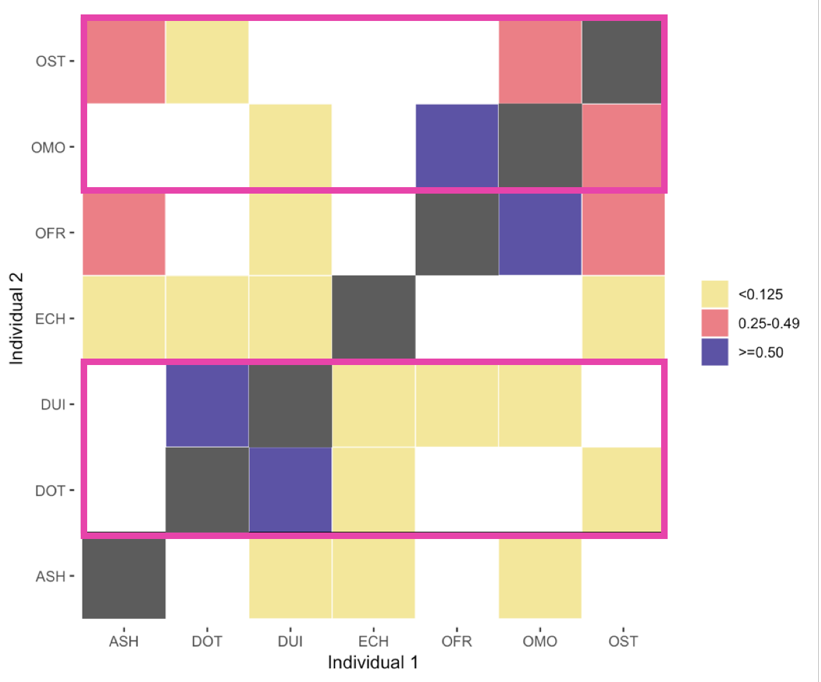

Figure S3. Maternal siblings have only partially overlapping grooming partners.

The matrix shows all of the adult females present in one small social group in one year, which represents a subset of our grooming dataset. Colored boxes represent pairs of individuals who groomed each other, and the color illustrates their pedigree relatedness; dark gray boxes on the diagonal would correspond to the individual self-grooming, a behavior that is not recorded. White boxes represent dyads who were never observed to groom in that year. Maternal siblings are grouped by pink outlines, illustrating that maternal siblings have only partially overlapping sets of grooming partners. It is not illustrated in this example, but our data also demonstrates that maternal siblings are not always equally related to a given grooming partner.

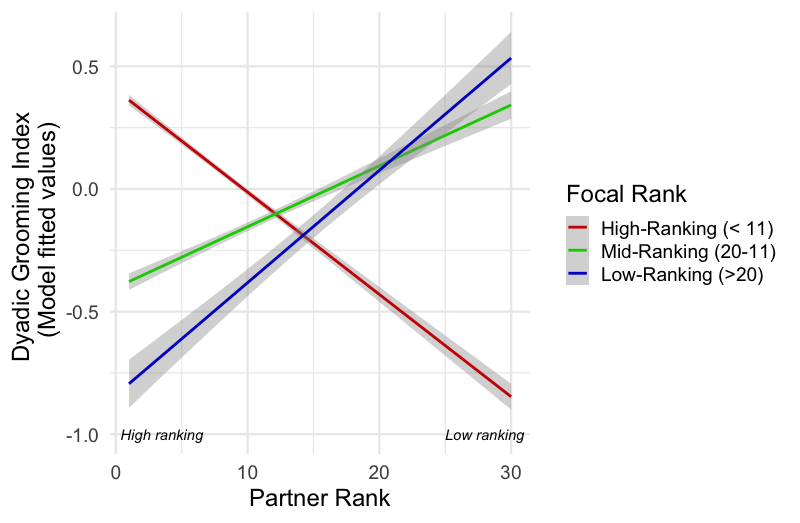

Figure S4. The dyadic grooming index is partially predicted by an interaction between the ordinal dominance ranks of the focal female and her partner.

High-ranking females (red line) gave more grooming to high-ranking partners (those with lower ordinal rank numbers), and the effect was in the opposite direction when focal females were middle or low-ranking (blue and green lines). Gray shaded areas represent 95% confidence intervals. Ranks are presented as categories for the purposes of visualization, but in the underlying model both focal and partner rank were modeled as continuous variables.

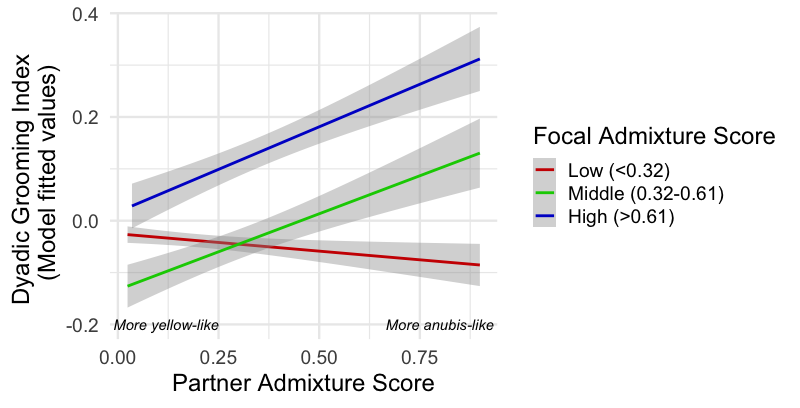

Figure S5. The dyadic grooming index is partially predicted by an interaction between focal and partner admixture score.

Focal females with low admixture scores (i.e., more yellow-like females; red line) gave approximately equal grooming to partners across the genetic ancestry spectrum. Individuals with more anubis ancestry (middle and high admixture scores) gave more grooming to partners with more anubis ancestry (blue and green lines). Gray shaded areas represent 95% confidence intervals. Admixture scores are presented as categories for the purposes of visualization, but in the underlying model both focal and partner admixture score were modeled as continuous variables.
